# The Evolution and Sequence Diversity of FhuA in *Salmonella* and *Escherichia*

**DOI:** 10.1101/320069

**Authors:** Yejun Wang, Xiongbin Chen, Guoqiang Zhu, Aaron P. White, Wolfgang Köster

**Author notes:** Corresponding authors: A.P.W., W.K.

## Abstract

The *fhuACDB* operon, present in a number of *Enterobacteriaceae*, encodes components essential for the uptake of ferric hydroxamate type siderophores. FhuA acts not only as transporter for physiologically important chelated ferric iron, but also as receptor for various bacteriophages, toxins and antibiotics, which are pathogenic to bacterial cells. In this research, the *fhuA* gene distribution and sequence diversity were investigated in *Enterobacteriaceae*, especially *Salmonella* and *Escherichia*. Comparative sequence analysis resulted in a *fhuA* phylogenetic tree that did not match the expected phylogeny based on housekeeping sequence analysis or trees of *fhuCDB* genes. The *fhuA* sequences showed a unique mosaic-clustering pattern. On the other hand, the gene sequences showed high conservation for strains from the same serovar or serotype. In total, six clusters were identified from FhuA proteins in *Salmonella* and *Escherichia*, among which typical peptide fragment variations could be defined. Six fragmental insertions / deletions and two substitution fragments were discovered, which could well classify the different clusters. Structure modeling demonstrated that all the six featured insertions/deletions and one substitution fragment are located at the apexes of the long loops of FhuA external pocket. These frequently mutated regions are likely under high selection pressure, and bacterial strains could have escaped from phage infection or toxin / antibiotics attack via *fhuA* gene mutations while maintaining the siderophore uptake activity essential for bacterial survival. The unusual *fhuA* clustering suggests that high frequency exchange of *fhuA* genes has occurred between enterobacterial strains after distinctive species were established.

**IMPORTANCE:** The enterobacterial *fhuACDB* operon encodes proteins which mediate the uptake of siderophores to supply the cells with iron essential for bacterial survival. Here we show different evolutionary patterns for the *fhu* genes within the same operon. The *fhuA* has a phylogenetic tree that does not match the species phylogeny, whereas the rest of the *fhu* genes do. The *fhuA* genes showed inter-species sequence convergence and conservation within specific serovars and serotypes. Nearly all of the significant sequence differences among FhuA clusters are located in potential ligand-binding sites on the extracellular surface of fhuA-encoding receptors. The unusual *fhuA* clustering suggests the frequent recombination and exchange of *fhuA* genes between enterobacterial strains in the evolutionary state after distinctive species were established.

Our findings suggested either a new evolutionary mechanism or local gene recombination in *fhuA* that is in contrast to previous evolutionary hypotheses that have formed under the assumption of no recombination.

## INTRODUCTION

FhuA (<u>f</u>erric <u>h</u>ydroxamate <u>u</u>ptake protein <u>A</u>) is a multi-functional protein, with a total length of 747 residues in *E. coli* strain MG1655, including an N-terminal 33 amino acid signal peptide and the 714 amino acid mature protein sequence (1). It represents a transmembrane receptor, residing in the outer membrane, which can mediate the uptake of ferric iron (bound to the siderophore ferrichrome) into the periplasm in a TonB-dependent manner (2, 3, 4). Iron is essential for bacterial survival and virulence, but it is mainly present in the non-soluble ferric form, and the bioavailable concentration in the environment is often extremely low. To increase bioavailability, bacteria secrete siderophore molecules that can chelate ferric iron with high affinity (5, 6). The iron-loaded siderophores can be recognized by bacterial membrane receptors (e.g., FhuA) and transported into periplasm. Further translocation of siderophores across the inner membrane is mediated by ABC transporters. In case of ferrichrome the transport into the cytoplasm depends on the binding protein FhuD, the integral membrane protein FhuB and the ATPase FhuC (7, and references therein). Besides the major physiological activity, FhuA is also usurped pathologically as the target for bacteriophages (T1, T5, ϕ80, and UC-1) and bacterial toxins (colicin M and microcin 25) (8). Some antibiotics also use FhuA as the receptor, e.g., albomycin and rifamycin CGP 4832 (9, 10).

Functional FhuA, embedded in the outer membrane, is a monomeric β-barrel protein, with its C-terminus forming 22 antiparallel β-strands and the N-terminus folded inside the β-barrel from the periplasmic side to form a cork domain (3, 11). The cork domain further separates a pair of pockets, the larger one open to the external medium and the smaller one facing the periplasm (3). The N-terminal portion of FhuA that contains the TonB box lies in the periplasm (4, 12, 13). The β-strands are connected by 11 long loops at the outer surface (L1~L11) and 10 short turns in the periplasmic side (T1~T10) (3, 11). The exact residues or motifs specifically interacting with ferrichrome within the external pocket of FhuA remain elusive, though it was found that some conserved residues in loops L3 and L11 were important for binding (3, 14). Ferrichrome binding induces amplified structural changes that go through the entire FhuA molecule up to the N-terminus, further signaling TonB binding and activating the TonB-ExbB-ExbD transporting system to internalize ferrichrome molecules (3, 15, 16). Other FhuA-binding molecules/ligands are also transported via the TonB-dependent pathway, except for phage T5 (15, 17). The binding sites and specificity determinants for these components are frequently located within the external loops of FhuA. For example, loops L4, L5 and L8 are involved in binding specificity of phages T1 and φ80, L5 and L8 are important for phage T5, and L3, L4, L7, L8 and L11 are involved in the sensitivity to colicin M and antibiotics albomycin and rifamycin (14, 15). Microcin J25 (MccJ25) binds at a similar location of FhuA as ferrichrome (18).

In *E. coli, fhuA* is located within an operon *fhuACDB*, comprising four genes in a conserved transcription direction of *fhuA, fhuC, fhuD* and *fhuB* (19). The protein products of *fhuC*, *fhuD* and *fhuB* form a complex, mediating the translocation of ferrichrome from the periplasm into the cytoplasm (20, 21, 22). However, in other species, *fhuA* and *fhuCDB* often appear in the same operon but with different transcription order, or within different operons (23, 24, 25). Fragmented gene sequences have been reported for *fhuA* frequently (26, 27). Gene evolutionary studies also suggested positive selection in *fhuA* rather than *fhuB, fhuC* or *fhuD* (28, 29). Therefore, FhuA may have a different evolutionary route with selection pressures that are independent from the rest of the FhuC/D/B system. Moreover, FhuD and FhuB in Gram negative bacteria display a broader ligand specificity compared to that of the OM receptor proteins (such as FhuA) in that they accept and transport not only ferrichromes but also other siderophores of the hydroxamate type, e.g. aerobactin and coprogen (20).

Despite the large amount of studies on *fhuA*, the mechanisms of the gene evolution and their relationship with protein function remain largely unknown. Nearly all studies have been limited to *fhuA* genes that are present in one or a few closely related strains. There is a lack of study on the gene distribution and sequence diversity among a large variety of bacterial strains. To partly address these points, in this research, we collected *fhuA* genes from a comprehensive list of *Salmonella* and *Escherichia* serovars and other close species, followed by observation of the gene distribution, sequence diversity and patterns and structure relevance. The resulting phylogenetic tree of *fhuA* clusters did not match the expected phylogeny based on housekeeping gene sequence analysis. The *fhuA* genes from various *Salmonella* serovars were found in distinct clusters; surprisingly together with *fhuA* genes from other species. This suggests an exchange of *fhuA* genes between different enterobacterial strains at an evolutionary state after distinctive species were established. In contrast, the *fhuCDB* genes located in the same operon downstream of *fhuA* as well as other TonB dependent receptor genes follow the commonly accepted phylogenic tree which clearly separates *Salmonella* strains from *Escherichia coli, Shigella, Citrobacter, Klebsiella* strains and other isolates.

## RESULTS

### 1. The distribution and diversification of *fhuA* in *Salmonella* and *Escherichia*

The representative genomes of *Salmonella* species, i.e., *S. enterica* and *S. bongori*, and six subspecies of *S. enterica*, including *enterica, arizonae, diarizonae, indica, salamae* and *houtenae* were screened, with a unique chromosomal copy of *fhuA* being identified in each strain. The order for *fhuA* and adjacent genes showed high conservation among *Salmonella* phylogenetic clusters (Fig 1a, b). Extended length of genomic sequences flanking *fhuA* showed apparent collinearity between different *Salmonella* genomes, further demonstrating the ancient presence of the *fhuA* locus in *Salmonella* (Fig 1c, upper). The *fhuA* locus even showed high gene synteny and sequence collinearity between *Salmonella* and *E. coli* strains, indicating its existence at least before the divergence of the two genera (Fig 1b; Fig 1c, lower).

**Fig 1.**
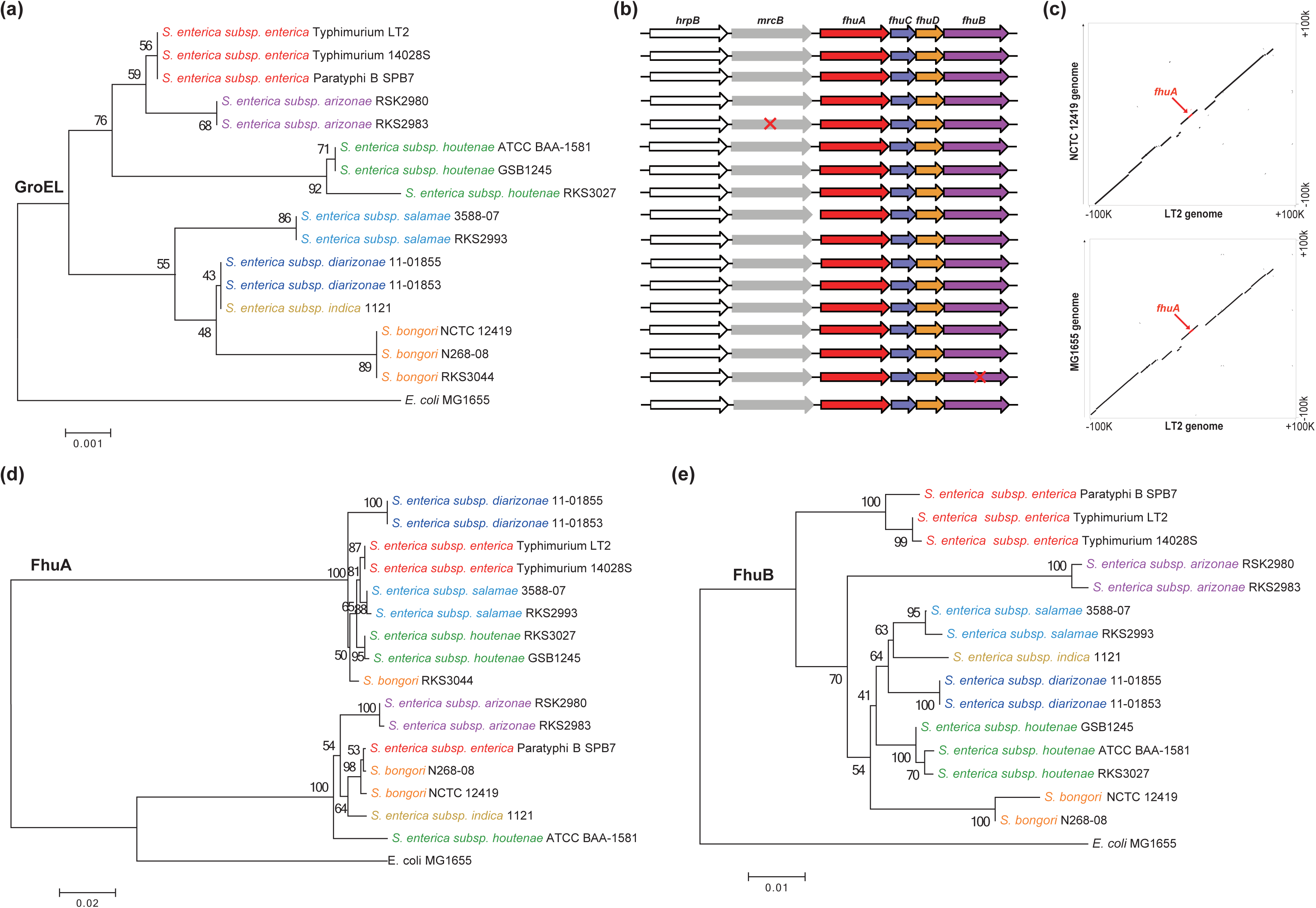
Distribution and phylogenetics of *fhuA* in *Salmonella*. (**a**) Neighbor-joining tree of GroEL, a protein encoded by the house-keeping gene *groEL*. Different species or subspecies are shown in different colors. (**b**) Gene order and distribution of *fhuA* locus in *Salmonella*. The strains were in line with those shown in the tree of (a). The pseudogene was indicated with ‘x’ in red. (**c**) Collinearity of fhuA locus between *S. bongori* NCTC 12419 (upper) or *E. coli* MG1655 (lower) and *S*. typhimurium LT2, respectively. (**d**)-(**e**) Neighbor-joining tree of FhuA and FhuB protein in *Salomonella*, respectively. *E. coli* MG1655 was used as the outgroup.

The phylogenetic tree was built for FhuA proteins among *Salmonella* strains (Fig 1d). Surprisingly, the topology was far different from the phylogenetic tree constructed based on a housekeeping gene, *groEL* (Fig 1a). Despite poor robustness, the GroEL tree should generally reflect the evolution of *Salmonella* species or subspecies, with strains clustered together within the same phylogenetic group (Fig 1a). In the FhuA tree, however, the strains belonging to a unique phylogenetic group diverged while the ones from different phylogenetic groups clustered (Fig 1d). In addition, the branch lengths between strains or species were very short, suggesting recent acquisition. The pattern was independent of the methods for tree reconstruction or sequence nature (Fig S1a, b). In contrast, patterns similar to the housekeeping gene tree were disclosed repeatedly for the trees based on other genes located in the same *fhuACDB* operon (Fig 1e; Fig S1c) or close to *fhuA* in the genome (Fig S1d).

The *fhuA* variance within a phylogenetic group was also confirmed by the protein tree reconstructed for the *Escherichia* genus, which included *E. fergausonii* and the *E. coli* / *Shigella* complex (Fig 2a). FhuA clustered into four clearly discernible groups, totally independent of phylogenetic clustering results of *Escherichia* / *Shigella* strains based on core gene datasets or oligonucleotide composition (Fig 2a) (30, 31). The FhuB protein tree could not form robust clusters due to the ultra-conservation of corresponding sequences (Fig S2). When the proteins from representative *Salmonella* and *Escherichia* strains were put together, the trees showed more striking difference between FhuA and FhuB. The FhuB tree was apparently consistent with the evolutionary trace of *Escherichia* and *Salmonella* species (Fig 2b), but the FhuA tree was clearly different (Fig 2c).

**Fig 2.**
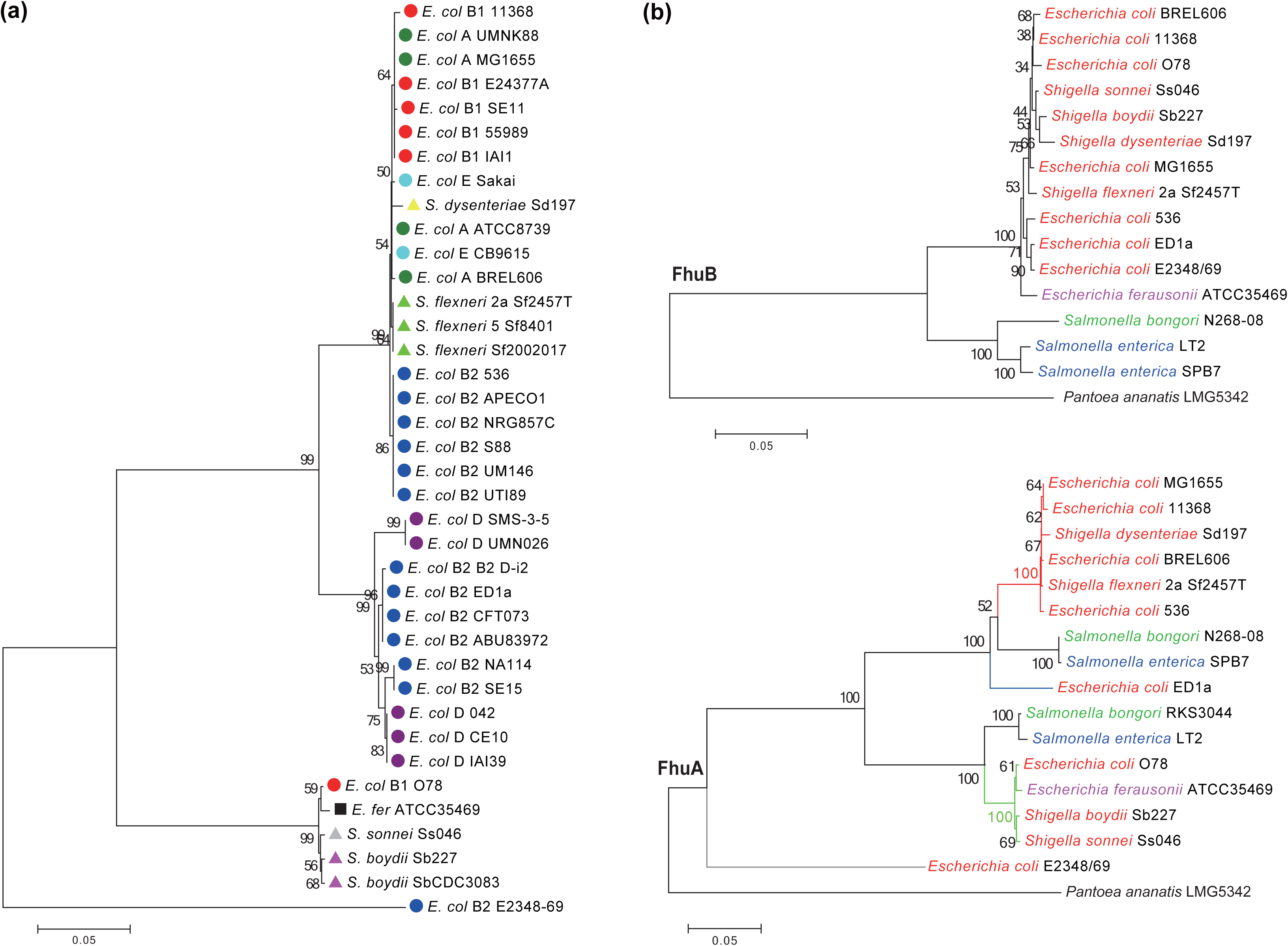
Phylogenetics of FhuA in *Escherichia* and *Salmonella*. (**a**) Neighbor-joining tree of FhuA among *E. coli* or *Shigella* strains. Different subgroups of *E. coli* / *Shigella* strains are shown in different colors or shapes. Neighbor-joining trees for FhuB (**b**) and FhuA (**c**) among *E. coli* / *Shigella* and *Salmonella* strains were constructed using *Pantoea ananatis* LMG5342 as the outgroup.

### 2. Within-serovar/serotype conservation of *fhuA* in *Salmonella* and *E. coli*

There were 112 serovars of *Salmonella enterica subsp. enterica* with at least one strain with a sequenced genome and 66 serovars with draft genomes for two or more strains (Table S1; http://www.ncbi.nlm.nih.gov/genome). Two or three representative genomes were selected randomly from each serovar with multiple genome-available strains, followed by screening and clustering analysis of the *fhuA* gene. No complete protein-encoding frame of *fhuA* was detected in strains from Typhi and Paratyphi A serovars, and a within-genome check indicated the existence but premature translation termination of *fhuA* gene in them (Table S1). For serovars Mississippi, Give, Javiana, Pulorum and Johannesburg, one strain from each serovar was detected with an incomplete fhuA-encoding frame (Table S1). The remaining 59 multi-strain serovars all contained a complete protein-encoding *fhuA* gene. An unsupervised clustering analysis demonstrated that the *Salmonella* FhuA proteins converged robustly into two clusters (S1 and S2) (Fig. 3a). For 54 of 59 serovars, the FhuA proteins from the same serovar clustered together, with serovars Bareilly, Mbandaka, Saintpaul, Senftenberg and Wandsworth being the exceptions (Fig 3a). The within-serovar conservation was not the result of selecting only a limited number of strains from each serovar, since inclusion of all sequenced genomes of Typhimurium (41 strains), Enteritidis (58 strains), Heidelberg (23 strains), Newport (19 strains) and Anatum (13 strains) led to similar S1 and S2 clustering (Fig. S3a; Table S1). Only two strains from these larger serovars did not match the observed pattern, Typhimurium FORC 015 and Newport CDC 2010K-2159 (Fig S3a; Table S1). Moreover, FhuA proteins from the serovars with a single strain whose genome was sequenced were also incorporated but no new pattern was disclosed (Fig S3b; Table S1).

**Fig 3.**
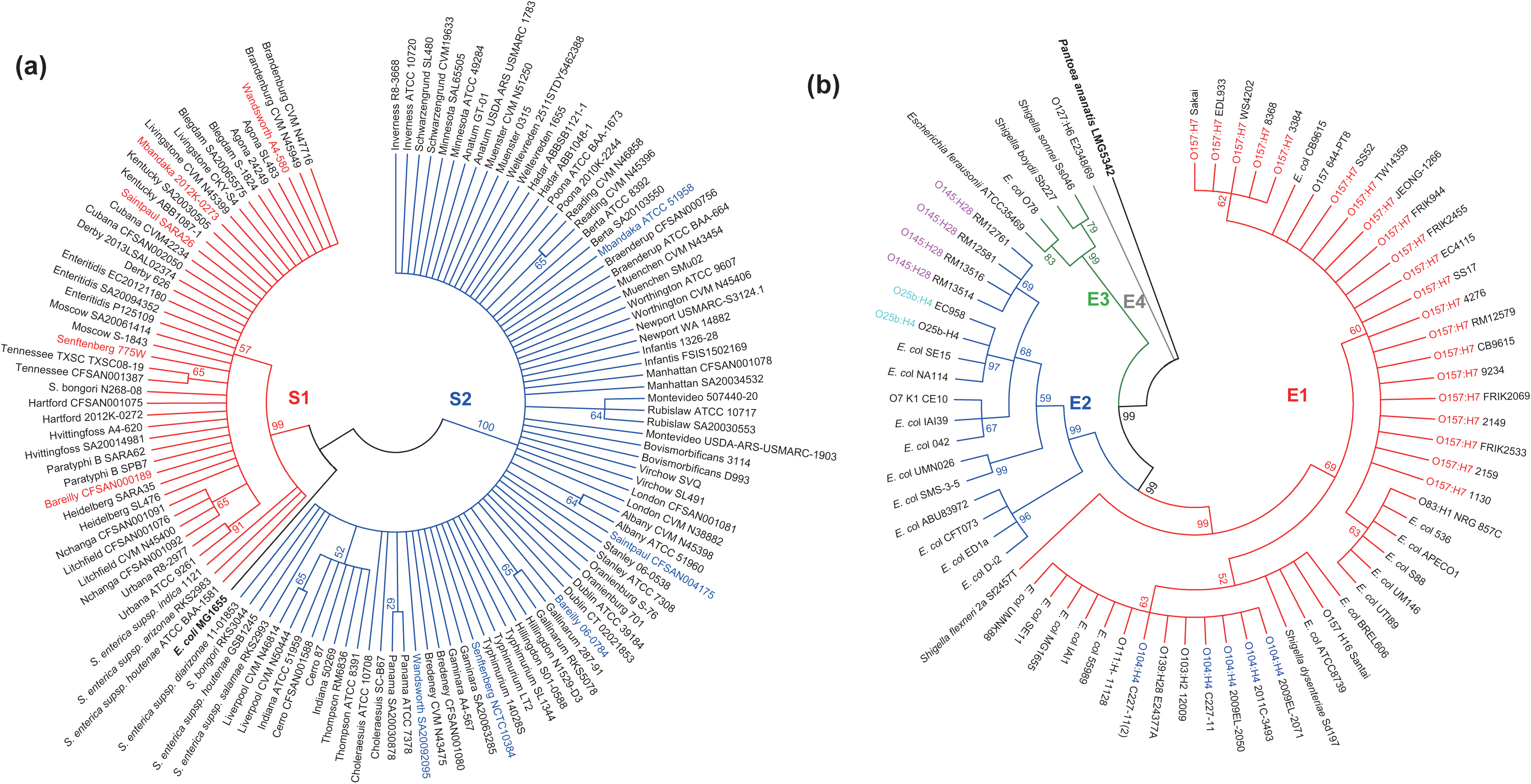
Distribution of FhuA clusters among serotypes of *Salmonella* and *E. coli*. (**a**) FhuA clusters in *Salmonella* serovars. The strains belonging to the same serovar but with different cluster of FhuA are shown in red or blue for their names. (**b**) FhuA clusters in *E. coli* serotypes.

Inclusion of more comprehensive *E. coli* (or *Shigella*) genomes available led to more definitive identification of the four major FhuA clusters (i.e., E1, E2, E3 and E4; Fig. 3b), as suggested previously (Fig. 2a). The FhuA proteins of multi-strain serotypes of *E. coli*, including O157:H7 (19), O145:H28 (4), O104:H4 (4) and O25b:H4 (2), also exhibited high within-serotype conservation (Fig 3b; Table S1).

### 3. Categorization of FhuA proteins with specific sequence features

FhuA protein sequences from the six clusters identified for *Salmonella* (S1 and S2) and *Escherichia* (E1-E4) were compared and determined to form three large groups, one with E1, S1 and E2, another with E3 and S2, and the third comprised of E4 solely (Fig. 4a). Sequence alignment also disclosed the insertions/deletions that featured each major group (E1/S1/E2, E3/S2, E4), subgroup (E1/S1, E2) or individual cluster (E1, S1, E3, S2) (Fig. 4b; Table S2). Group E4 proteins had unique insertions in two positions termed Indel 1 (residues 297 ~ 300) and Indel 2 (residues 699 ~ 703) (Fig 4b). There was also a divergent fragment insertion in the same regions of *Pantoea* FhuA, which served as the outgroup for the phylogenetic tree (Fig 4b). In contrast, the E1/S1/E2 FhuA proteins all had peptides inserted in two different regions termed Indel 3 (residues 361 ~ 383) and Indel 4 (residues 459 ~ 465), with apparent sequence homology between E1 and S1 rather than E2 (Fig 4c). For the E3/S2 group, unique peptide insertions were detected at Indel 5 (residues 517 ~ 521) and Indel 6 (residues 564 ~ 568) (Fig 4b). The E1 and S1 (or E3 and S2) FhuA proteins had high similarity and no unique insertions/deletions could be detected between them. Amino acid substitutions were sufficient to distinguish each individual group apart, however, since the insertions/deletions were removed before constructing the clustering tree. The distinct amino acid sequences were detected at regions termed Sub1 (residues 415 – 417) between E3 and S2 and Sub2 (residues 242 – 244) between E1 and S1 (Fig 4b).

**Fig 4.**
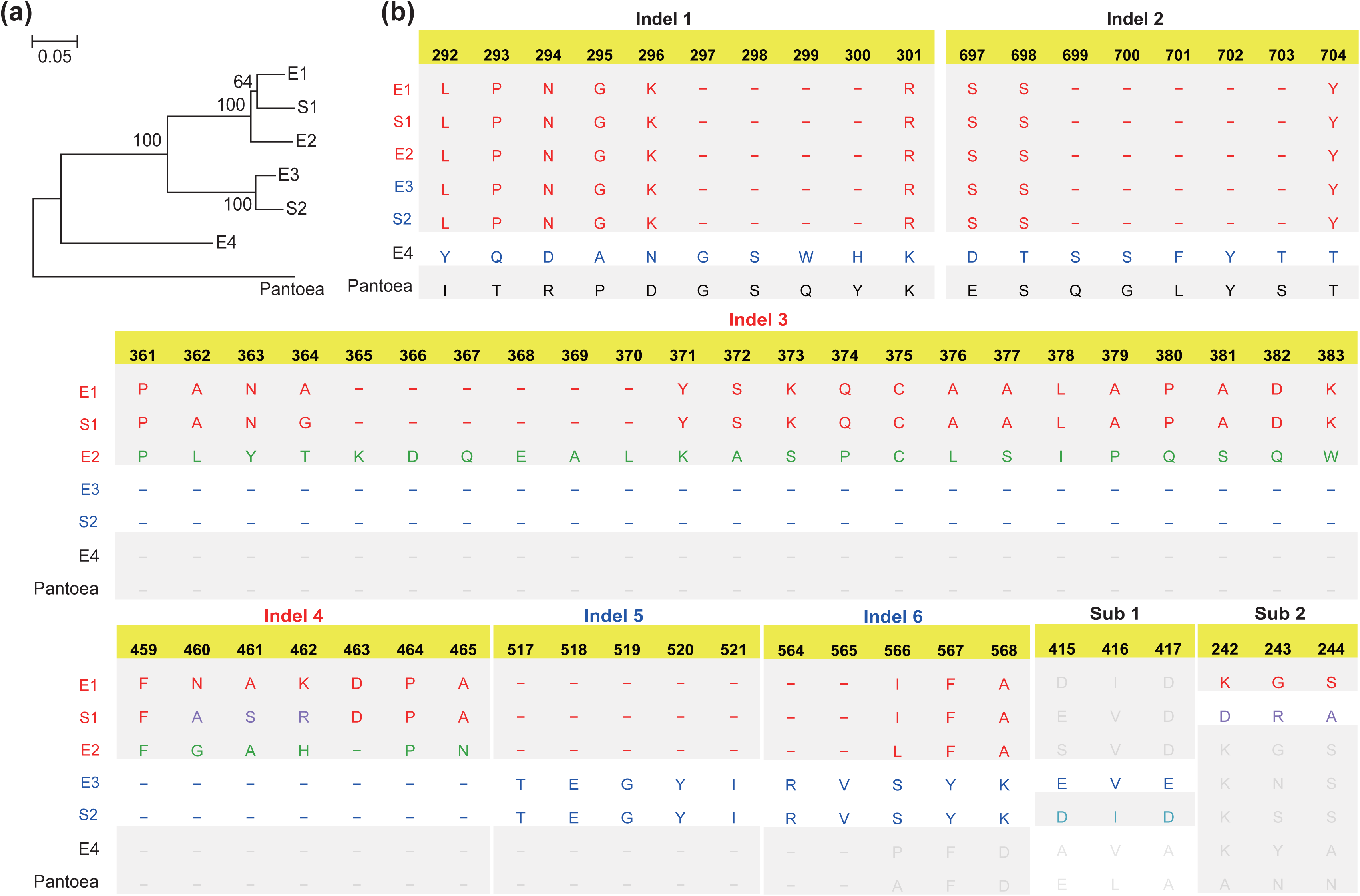
Typical mutation patterns of FhuA clusters in *Salmonella* and *Escherichia*. (**a**) A diagram tree showing the clusters of FhuA in *Salmonella* and *Escherichia*. (**b**) The typical indels or substitutions that can distinguish different FhuA clusters. Indel 1 and Indel 2 can distinguish E4 from E1/S1/E2/E3/S2 (**b**), Indel 3-6 can distinguish E3/S2 from E1/S1/E2; Sub 1 can distinguish E3 from S2; and Sub2 can distinguish S1 from E1.

To enable a more thorough categorization for FhuA, more representative genomes of *Enterobacteriaceae* were screened for possible *fhuA* genes. Besides *Escherichia*, *Salmonella* and *Pantoea*, *fhuA* was detected in *Cronobacter*, *Enterobacter, Erwinia, Klebsiella, Rahnella, Raoultella* and *Serratia* (Fig 5a). With the inclusion of sequences from these new organisms, the FhuA proteins were clustered into four major groups (I ~ IV), where E1/S1/E2 and E3/S2 belonged to Group I, E4 belonged to Group II and *Pantoea* FhuA belonged to Group IV (Fig 5a). Interestingly, the main variation for insertions/deletions among the four major groups remained in the six locations identified previously (i.e., Indel 1 ~ 6), suggesting the easy gain or loss and variation of fragments in these positions (Fig 5b). The FhuA N-terminal region of approximately 50 amino acids was also highly variable, which represents the signal peptide sequences and the unstructured fragment (3, 11). However, the TonB-box was conserved (Fig S4). Sequences between N-terminal ~50 and 200 amino acids were more conserved than other regions. FhuA sequences with all the Indels and flanking variable amino acid positions removed or N/C-terminal fragments truncated for the N1-200 amino acids retained similar clustering patterns with original FhuA proteins, indicating the group-specific variations were not constrained to the insertion/deletion loci or specific fragments (Fig 5c).

**Fig 5.**
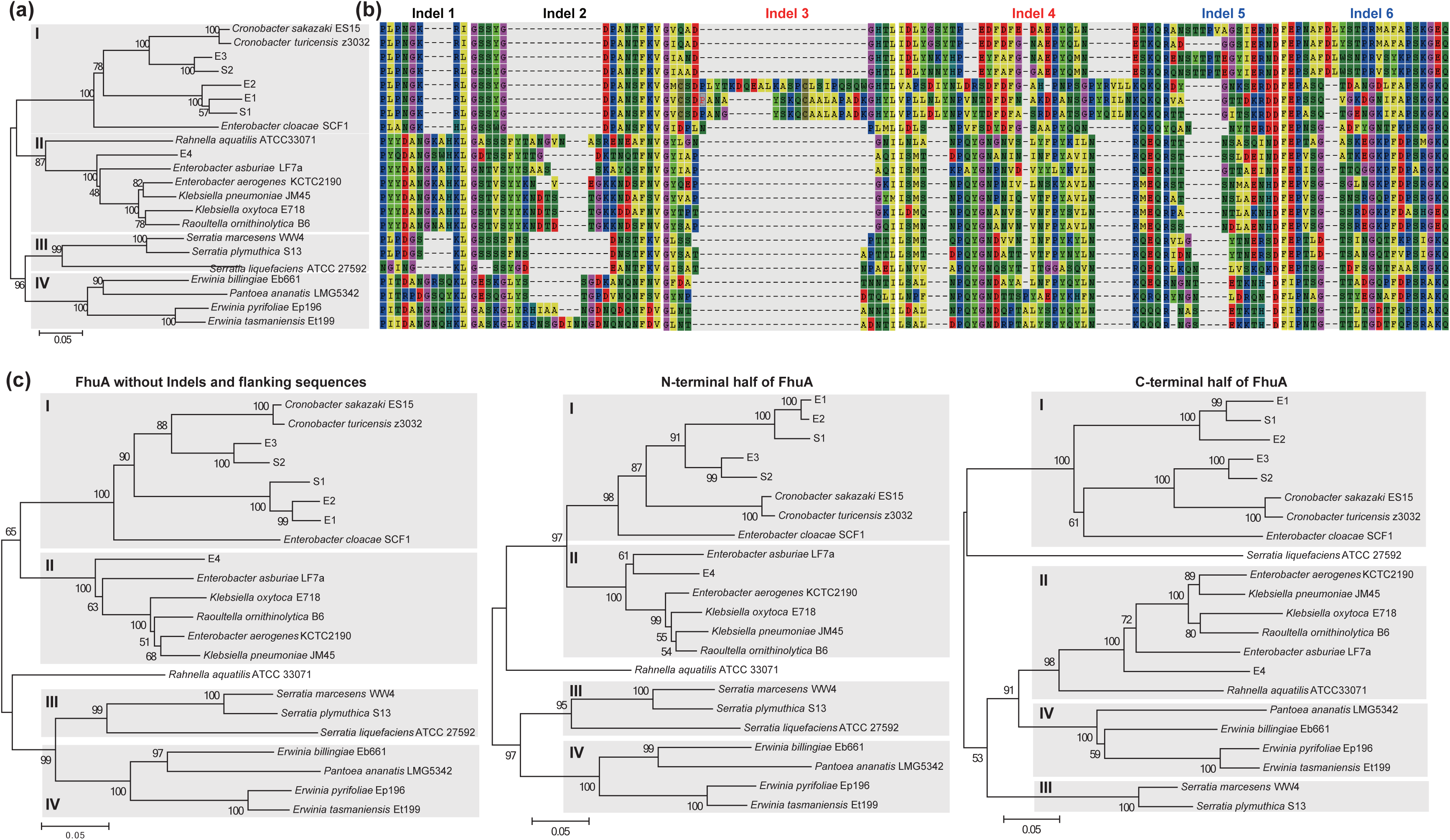
Typical mutation patterns of FhuA clusters in *Enterobacteriaceae*. (**a**) Neighbor-joining tree of FhuA protein in *Enterobacteriaceae* strains. The tree grouped FhuA into 4 clusters (I ~ IV). (**b**) The indel patterns of different FhuA clusters. (**c**) Neighbor-joining tree of FhuA proteins after removing the Indels and flanking highly mutated sequences (left), N-terminal (middle) and C-terminal half (right) FhuA protein sequences, respectively.

### 4. Structure variations in the extracellular surface of FhuA proteins

The atomic structure of FhuA from *E. coli* K-12 (group E1) was solved (3, 11). Based on the coordinates of the structure, we performed homology modeling to compute the structures of FhuA proteins from groups E2, E3, E4, S1, S2, as well as from *Pantoea* and *Serratia*. Generally, all the proteins had a similar, predicted cork-barrel structure, with cork domains showing high similarity in both sequence and structure (Fig 6). Interestingly, the six Indel regions previously identified were located in loops of the external pocket of FhuA that is known to face the extracellular environment, corresponding to loops L3 (Indel 1), L10 (Indel 2), L4 (Indel 3), L5 (Indel 4), L6 (Indel 5), and L7 (Indel 6) (Fig 6a; Table S2). One of the key substitution regions that helped to differentiate FhuA into subgroups also happened in an external surface loop - L2 (Sub 2), whereas the other substitution region located in a linker between β -strands (T4 for Sub 1) (Fig 6a). Few variations were observed in either cork or transmembrane barrels, or even the surface facing the periplasm, coinciding with the results of sequence alignment (Table S2).

**Fig 6.**
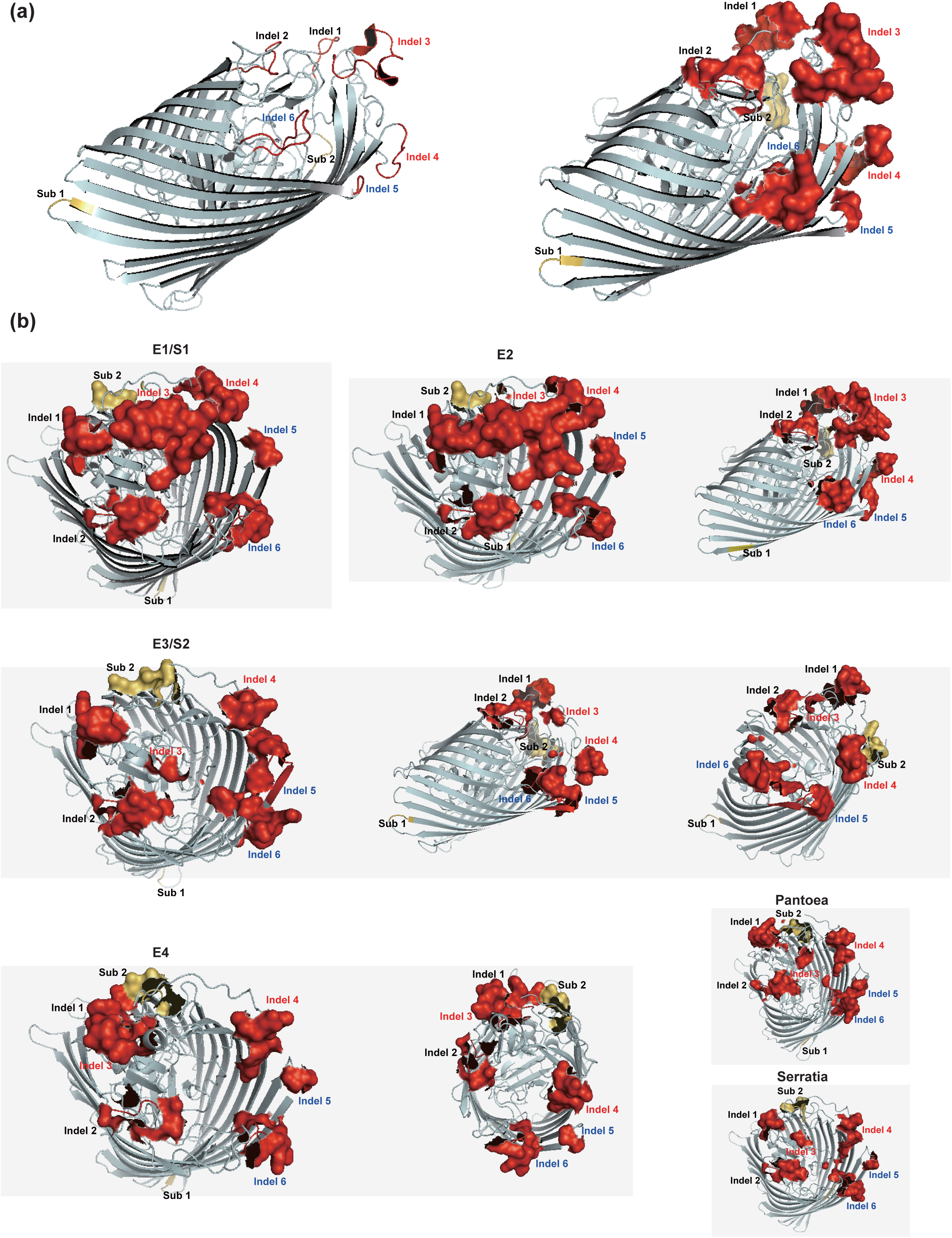
Influence of typical mutations on FhuA structure. (**a**) Resolved E. coli FhuA structure (PDB accession: 1QFF) with mutation hotspots highlighted. (**b**) Modeling structure of different FhuA clusters (E1/S1, E2, E3/S2, E4, Pantoea, Serratia), with mutation hotspots highlighted. The overall protein structure was displayed with PyMOL in cartoon format; the mutation hostspots were shown in either cartoon or surface format.Table S1. FhuA patterns in *Salmonella* and *E. coli* strains.

S1 FhuA proteins had a predicted structure that was highly similar to E1, while E2 proteins showed an insertion in L4 (Indel 3) when compared to E1/S1, causing the approaching of the domain with L3 (Indel 1) and the formation of a compact surface structure (Fig 6B, E2, left and right). In E3 proteins, L4 was shortened extensively, along with L5, but both L6 and L7 were elongated and formed a fused surface (Fig 6b, E3, left, middle and right). E4 proteins showed a more diverse external surface structure, which had an insertion in L3 and L10, a deletion in L4 and striking changes in loops L2, L5, L6 and L7. The loops L2 (Sub2), L3 (Indel 1) and L4 (Indel 2) became continuous forming a local surface structure (Fig 6B, E4, left and right). The *Pantoea* and *Serratia* FhuAs, also showed diverse outer surface structure but conservation in other regions (Fig 6B, Pantoea and Serratia).

Taken together, the structure modeling analysis suggested that the diversity of FhuA sequences was represented in the structural variation in the exposed regions facing the external environment.

## DISCUSSION

The *fhuACDB* locus showed high conservation within the genomes of *Salmonella* and *Escherichia*. Despite the fragmentation of the *fhuA* protein-coding frame in some serovars or strains, e.g., *Salmonella* serovar Typhi, the gene vestige can be well traced like the other genes within the same operon, i.e., *fhuC, fhuD* and *fhuB*. This indicates that *fhuA* was not likely acquired via horizontal gene transfer within the strains after the divergence of *Salmonella* and *Escherichia*. However, the mosaic *fhuA* gene and protein sequences do not follow the phylogenetic routes of species. This contrasts the findings resulting from comparing *fhuC/D/B* or housekeeping genes among *Salmonella* or *E. coli* strains, which shows a clear separation of those species. There have been previous reports that *fhuA* is subject to positive selection, which may explain the sequence variation. For example, Chen et al firstly reported positive selection of *fhuA* gene specifically in uropathogenic *E. coli* (UPEC) (28). Petersen et al. further found that the gene was positively selected among multiple *E. coli* strains (29). We also found within-serovar/serotype conservation and between-serovar/serotype diversity of *fhuA* sequences in either *Salmonella* or *E. coli*, which was consistent with the positive selection hypothesis within a single genus: the *fhuA* gene should vary to be better adapted to the specific environment where the serovars/serotype survives. However, a simple positive selection model has difficulty in explaining the inter-genus/species shared *fhuA* sequence clusters and mutation patterns disclosed in this study. Within a single genus or species, the ancient *fhuA* may have diversified as the strains evolved and adapted to different growth environments. The inter-genus/species common sequence clusters and mutation patterns could indicate convergent evolution of *fhuA* for strains living in similar environments but originating from different genera or species. It suggests for *fhuA* that the selection pressures based on the specific environment are stronger than taxon-conservation.

Sequence diversion of *fhuA* and the inter-genus/species common patterns observed are more reflected by short-fragment insertions/deletions than by single-nucleotide polymorphisms (29). Common insertions/deletions among widely varied gene sequences from different genera or species are more likely formed via sequence recombination. However, in *dn*/*ds* based selection studies, these inserted/deleted sites are removed before further analysis (32, 33). Moreover, the selection analysis on the *fhuA* gene was based on the premise of ‘no recombination happening in the gene’ (28, 29). Recombination events could have been missed by existent algorithms, especially when the number and variety of analyzed strain samples were limited, such as in the research of Chen et al. (2006) and Petersen et al. (2007). A more simplified explanation for the mosaic *fhuA* sequences observed is that frequent recombination events have shaped the *fhuA* sequence patterns. The amino acid substitution patterns within different peptide fragments of FhuA can classify the protein sequences into clusters similar to those grouped by insertion/deletion patterns. Therefore, we hypothesize that recombination has occurred throughout the entire *fhuA* gene.

Two distinct clusters for *fhuA* genes in *Salmonella* (S1 and S2) and four distinct clusters for *fhuA* genes in *E. coli* (E1~E4) were clearly defined in our study. S1 *fhuA* genes are more similar to E1 and E2, S2 *fhuA* genes are more similar to E3, and the E4 *fhuA* genes appear to be more independent. When the taxa were enlarged to include more Enterobacterial species, four *fhuA* super clusters (I~IV) were detected. We identified eight clear mutation hotspots in the *fhuA* protein sequence, corresponding to six insertion/deletion sites and two amino acid substitution fragments that mutated frequently within all the *Enterobacteriaceae* strains. Perhaps most interesting was that each Indel region and one of the substitution regions mapped to the apexes of FhuA surface loops (Fig 6). In contrast, the TonB box, β -strands and the cork showed high sequence conservation. Consequently, the predicted overall structure did not vary greatly between FhuA clusters, whereas the size and composition of outside loops varied a lot (Fig 6). The surface regions, especially the loops, are critical for recognition of various substrates, and determine the binding specificity (15, 14). It needs to be verified experimentally whether the different clusters of FhuA with loop conformation changes show different binding patterns to a variety of substrates, including bacteriophages. Selection pressures based on bacteriophage binding have been observed in the past (34, 35) and we hypothesize that evolved resistance to key bacteriophages could be responsible for some of the group-specific changes in surface-exposed FhuA loops. We recently found that *Yersinia* leucine-rich repeat genes had frequent mutations that were constrained to maintain protein activity and overall structure (36). Instead of random nucleotide substitutions, the amino acid codons tended to mutate with maximum parsimony (i.e., in the most straightforward and simplest way). Therefore, the observed mutations at the peptide level often showed predictable patterns rather than total irregularity (36). It is possible for a protein to have multiple activities, some of which may be conserved and essential for organism survival, and therefore under negative selection, while others could be placed under positive selection. Such proteins could adopt co-mutations at multiple loci to simultaneously change some function, such as alteration of a ligand-recognition motif, while still maintaining the essential activity (36). Under these two hypotheses, the mosaic evolution patterns, the mutation loci and the possible protein activities of FhuA could be well explained and associated.

There are many more interesting but unanswered questions related with *fhuA* evolution. For example, the *fhuA* and *fhuC/fhuD/fhuB* genes, located in the same genomic region, show the same gene order, and were reported to form an operon sharing uniform gene regulation in *E coli* (or *Salmonella)* strains (19, 21). However, in other species, the gene order could be disturbed, and the genes could be split into different operons (23, 24, 25). How did the expression regulation of *fhuA* evolve, and what is its relationship with *fhuC/D/B*? We only inspected the sequence patterns of *fhuA* in *Enterobacteriaceae*, and the distribution and evolution of *fhuA* beyond the family are yet largely unknown. Within a bacterial genome, generally very few genes (~30) are thought to be shaped by positive selection pressure (28, 29). Besides FhuA, other β -barrel porins, e.g., OmpF, OmpC and LamB, and iron acquisition related proteins, e.g., FepE, EntD and EntF, were also enriched in the list of proteins undergoing positive selection (28, 29). Such extraordinary enrichment of proteins with similar structure or function suggests the significance of positive selection on these proteins. A thorough analysis on each individual protein and a comparison could possibly inspire important insights into the evolution and function of these proteins. Finally, *fhuA* was annotated as ‘pseudogene’ in many strains (26, 27). In the case of S. Typhi *fhuA* it appears that the “pseudogene” is the result of small (one or two base pairs) deletions or insertions leading to a frame shift. Most likely, the loss of a functional *fhuA* gene could be tolerated in this human adapted serovar, which has less reliance on long-term survival in the environment where acquisition of ferric iron via ferrichrome would constitute a major fitness advantage. Most frame shift events in *fhuA* result in a significant polar effect regarding the expression of the downstream located *fhuCDB* genes. Polar *fhuA* mutants can be tolerated in Typhi but probably not in other serovars which would depend (at least in certain periods of their life cycle) on the uptake of various hydroxamate siderophores (such as aerobactin, coprogen, rhodotorulic acid etc.) transported by the FhuCDB complex

In summary, the unusual FhuA mosaic clustering suggests an exchange of *fhuA* genes between different enterobacterial strains at an evolutionary state after distinctive species such as *Escherichia* and *Salmonella* were established. For the moment, the “driving force” (i.e., selective pressure, phages, nutritional requirements) and mechanism(s) of genetic exchange (e.g., conjugation, transduction, mobile elements) remain largely unknown. It is also unclear for which period of time this genetic exchange was happening, and if this possibility still exists. It appears that there was no significant *fhuA* gene exchange between *Escherichia* and *Salmonella* and between different serovars in the near past since the FhuA protein shows invariance among strains belonging to the same *Salmonella* serovar or *E. coli* serotype. In general, all strains of a given serovar carry the same FhuA cluster type irrespective of their “lifestyle”, environment or geographical location.

## MATERIALS AND METHODS

### 1. Bacterial strains, genomes and gene sequences

The genomes of *Salmonella*, *Escherichia* and other *Enterobacteriaceae* strains were downloaded from the National Center for Biotechnology Information (NCBI) genome database. For each strain with a publicly available genome, the strain information, including serovar/serotype, taxonomy and pathogenicity, was manually annotated from the corresponding genome annotation files, original publications and NCBI taxonomy database. Genome annotation files were the base for retrieving gene and protein sequence. Standalone BLAST suite was downloaded from the NCBI website, installed, and used for screening FhuA, FhuB, FhuC, FhuD, GroEL and HrpB with *E. coli* MG1655 counterparts as the query sequences and with an identity cutoff of 70%. The genomic loci of the genes were positioned based on the protein description within corresponding genome annotation file. For strains without a specific protein captured, the potential genomic locus was reasoned according to its adjacent conserved genes, followed by retrieving, alignment and comparison of the nucleotide sequences.

### 2. Comparative genomics and phylogenetic analysis

The *fhuA* and neighbor genes were depicted according to their order present in their corresponding genome, to make synteny comparison among different bacterial strains. The interesting genes and their flanking sequences (100-kb each side) were also retrieved from the corresponding genomes, and PipMaker was used for collinearity analysis (37). The proteins or genes were aligned and phylogenetic analysis was performed with MEGA6.0 (38). Maximum likelihood or neighbor-joining method was adopted, with bootstrapping tests for 1,000 replicates.

### 3. Structure modeling and comparison

The experimentally resolved structure of *E. coli* FhuA was used as reference (PDB accession: 1QFF), and the structures of other FhuA proteins were predicted with PHYRE2 (39). PyMOL was used for structure visualization, analysis and comparison (http://www.pymol.org).

## Acknowledgement

The research was supported by a Natural Science Fund of Shenzhen (JCYJ201607115221141) and a Shenzhen Peacock Start-up Research Fund for Overseas Talents to YW.

## Supplementary Materials

**Table S1. FhuA patterns in *Salmonella* and *E. coli* strains.**

**Table S2. Protein mutation profiles distinguishing different FhuA clusters.**

**Fig S1. Different evolutionary routes of FhuA with other proteins in *Salmonella*.** (**a**) Maximum likelihood tree of FhuA protein. (**b**) Neighbor-joining tree of *fhuA* genes. (**c**)-(**d**) Neighbor-joining tree of FhuC and HrpB protein, respectively. The corresponding ortholog in *E. coli* MG1655 was used as the outgroup for each tree.

**Fig S2. Neighbor-joining tree of FhuB in *E. coli* / *Shigella* complex.** Different subgroups of *E. coli* / *Shigella* strains were shown in different colors or shapes. Boostrapping test score was indicated for each node.

**Fig S3. Distribution of FhuA clusters among *Salmonella* serovars.** (**a**) Distribution of FhuA clusters (**a**) among the multiple strains of Enteritidis and Typhimurium, or (**b**) among an enlarged number of serotypes.

**Fig S4. Conservation of TonB box in FhuA proteins from *Enterobacteria*. The box was highlighted with grey background in each sequence.**

## Reference

1. Coulton JW, Mason P, Cameron DR, Carmel G, Jean R, Rode HN. (1986). Protein fusions of beta-galactosidase to the ferrichrome-iron receptor of Escherichia coli K-12. J Bacteriol. 165(1):181–92.

2. Hantke K, Braun V. (1975). Membrane receptor dependent iron transport in Escherichia coli. FEBS Lett. 49(3):301–5.

3. Ferguson AD, Hofmann E, Coulton JW, Diederichs K, Welte W. (1998). Siderophore-mediated iron transport: crystal structure of FhuA with bound lipopolysaccharide. Science. 282(5397):2215–20.

4. Freed DM, Lukasik SM, Sikora A, Mokdad A, Cafiso DS. (2013). Monomeric TonB and the Ton box are required for the formation of a high-affinity transporter-TonB complex. Biochemistry. 52(15):2638–48.

5. Holden VI, Bachman MA. (2015). Diverging roles of bacterial siderophores during infection. Metallomics. 7(6):986–95.

6. Wilson BR, Bogdan AR, Miyazawa M, Hashimoto K, Tsuji Y. (2016). Siderophores in Iron Metabolism: From Mechanism to Therapy Potential. Trends Mol Med. 22(12):1077–1090.

7. Köster W. (2005). Cytoplasmic membrane iron permease systems in the bacterial cell envelope. Front Biosci. 10: 462–477.

8. Braun V. (2009). FhuA (TonA), the career of a protein. J Bacteriol. 191(11):3431–6.

9. Ferguson AD, Braun V, Fiedler HP, Coulton JW, Diederichs K, Welte W. (2000). Crystal structure of the antibiotic albomycin in complex with the outer membrane transporter FhuA. Protein Sci. 9(5):956–63.

10. Ferguson AD, Ködding J, Walker G, Bös C, Coulton JW, Diederichs K, Braun V, Welte W. (2001). Active transport of an antibiotic rifamycin derivative by the outer-membrane protein FhuA. Structure. 9(8):707–16.

11. Locher KP, Rees B, Koebnik R, Mitschler A, Moulinier L, Rosenbusch JP, Moras D. (1998). Transmembrane signaling across the ligand-gated FhuA receptor: crystal structures of free and ferrichrome-bound states reveal allosteric changes. Cell. 95(6):771–8.

12. Schöffler H, Braun V. (1989). Transport across the outer membrane of Escherichia coli K12 via the FhuA receptor is regulated by the TonB protein of the cytoplasmic membrane. Mol Gen Genet. 217(2–3):378–83.

13. Noinaj N, Guillier M, Barnard TJ, Buchanan SK. (2010). TonB-dependent transporters: regulation, structure, and function. Annu Rev Microbiol. 64: 43–60.

14. Endriss F, Braun V. (2004). Loop deletions indicate regions important for FhuA transport and receptor functions in Escherichia coli. J Bacteriol. 186(14):4818–23.

15. Moeck GS, Coulton JW. (1998). TonB-dependent iron acquisition: mechanisms of siderophore-mediated active transport. Mol Microbiol. 28(4):675–81.

16. Hickman SJ, Cooper REM, Bellucci L, Paci E, Brockwell DJ. (2017). Gating of TonB-dependent transporters by substrate-specific forced remodelling. Nat Commun. 8: 14804.

17. Flayhan A, Wien F, Paternostre M, Boulanger P, Breyton C. (2012). New insights into pb5, the receptor binding protein of bacteriophage T5, and its interaction with its Escherichia coli receptor FhuA. Biochimie. 94(9):1982–9.

18. Mathavan I, Zirah S, Mehmood S, Choudhury HG, Goulard C, Li Y, Robinson CV, Rebuffat S, Beis K. (2014). Structural basis for hijacking siderophore receptors by antimicrobial lasso peptides. Nat Chem Biol. 10(5):340–2.

19. Fecker L, Braun V. (1983). Cloning and expression of the fhu genes involved in iron(III)-hydroxamate uptake by Escherichia coli. J Bacteriol. 156(3):1301–14.

20. Köster W. (1991). Iron(III)hydroxamate transport across the cytoplasmic membrane of *Escherichia coli*. Biol Metals. 4: 23–32.

21. Mademidis A, Köster W. (1998). Transport activity of FhuA, FhuC, FhuD, and FhuB derivatives in a system free of polar effects, and stoichiometry of components involved in ferrichrome uptake. Mol Gen Genet. 258(1–2):156–65.

22. Carter DM, Miousse IR, Gagnon JN, Martinez E, Clements A, Lee J, Hancock MA, Gagnon H, Pawelek PD, Coulton JW. (2006). Interactions between TonB from Escherichia coli and the periplasmic protein FhuD. J Biol Chem. 281(46):35413–24.

23. Mikael LG, Pawelek PD, Labrie J, Sirois M, Coulton JW, Jacques M. (2002). Molecular cloning and characterization of the ferric hydroxamate uptake (fhu) operon in Actinobacillus pleuropneumoniae. Microbiology. 148(Pt 9):2869–82.

24. del Río ML, Navas J, Martín AJ, Gutiérrez CB, Rodríguez-Barbosa JI, Rodríguez Ferri EF. (2006). Molecular characterization of Haemophilus parasuis ferric hydroxamate uptake (fhu) genes and constitutive expression of the FhuA receptor. Vet Res. 37(1):49–59.

25. Abdelhamed H, Lu J, Lawrence ML, Karsi A. (2016). Ferric hydroxamate uptake system contributes to Edwardsiella ictaluri virulence. Microb Pathog. 100: 195–200.

26. Stevens JB, Carter RA, Hussain H, Carson KC, Dilworth MJ, Johnston AW. (1999). The fhu genes of Rhizobium leguminosarum, specifying siderophore uptake proteins: fhuDCB are adjacent to a pseudogene version of fhuA. Microbiology. 145 (Pt 3):593–601.

27. Parkhill J, Dougan G, James KD, Thomson NR, Pickard D, Wain J, Churcher C, Mungall KL, Bentley SD, Holden MT, Sebaihia M, Baker S, Basham D, Brooks K, Chillingworth T, Connerton P, Cronin A, Davis P, Davies RM, Dowd L, White N, Farrar J, Feltwell T, Hamlin N, Haque A, Hien TT, Holroyd S, Jagels K, Krogh A, Larsen TS, Leather S, Moule S, O'Gaora P, Parry C, Quail M, Rutherford K, Simmonds M, Skelton J, Stevens K, Whitehead S, Barrell BG. (2001). Complete genome sequence of a multiple drug resistant Salmonella enterica serovar Typhi CT18. Nature. 413(6858):848–52.

28. Chen SL, Hung CS, Xu J, Reigstad CS, Magrini V, Sabo A, Blasiar D, Bieri T, Meyer RR, Ozersky P, Armstrong JR, Fulton RS, Latreille JP, Spieth J, Hooton TM, Mardis ER, Hultgren SJ, Gordon JI. (2006). Identification of genes subject to positive selection in uropathogenic strains of Escherichia coli: a comparative genomics approach. Proc Natl Acad Sci U S A. 103(15):5977–82.

29. Petersen L, Bollback JP, Dimmic M, Hubisz M, Nielsen R. (2007). Genes under positive selection in Escherichia coli. Genome Res. 17(9):1336–43.

30. Touchon M, Hoede C, Tenaillon O, Barbe V, Baeriswyl S, Bidet P, Bingen E, Bonacorsi S, Bouchier C, Bouvet O, Calteau A, Chiapello H, Clermont O, Cruveiller S, Danchin A, Diard M, Dossat C, Karoui ME, Frapy E, Garry L, Ghigo JM, Gilles AM, Johnson J, Le Bouguénec C, Lescat M, Mangenot S, Martinez-Jéhanne V, Matic I, Nassif X, Oztas S, Petit MA, Pichon C, Rouy Z, Ruf CS, Schneider D, Tourret J, Vacherie B, Vallenet D, Médigue C, Rocha EP, Denamur E. (2009). Organised genome dynamics in the Escherichia coli species results in highly diverse adaptive paths. PLoS Genet. 5(1):e1000344.

31. Sims GE, Kim SH. (2011). Whole-genome phylogeny of Escherichia coli/Shigella group by feature frequency profiles (FFPs). Proc Natl Acad Sci USA. 108(20):8329–34.

32. Hughes AL, Nei M. (1988). Pattern of nucleotide substitution at major histocompatibility complex class I loci reveals overdominant selection. Nature. 335: 167–170.

33. Yang Z, Bielawski JP. (2000). Statistical methods for detecting molecular adaptation. Trends Ecol. Evol. 15: 496–503.

34. Seed KD, Yen M, Shapiro BJ, Hilaire IJ, Charles RC, Teng JE, Ivers LC, Boncy J, Harris JB, Camilli A. (2014). Evolutionary consequences of intra-patient phage predation on microbial populations. eLife. 3:e03497.

35. van Houte S, Buckling A, Westra ER. (2016). Evolutionary Ecology of Prokaryotic Immune Mechanisms. Microbiol Mol Biol Rev. 80(3):745–63.

36. Hu Y, Huang H, Hui X, Cheng X, White AP, Zhao Z, Wang Y. (2016). Distribution and Evolution of Yersinia Leucine-Rich Repeat Proteins. Infect Immun. 84(8):2243–2254.

37. Schwartz S, Zhang Z, Frazer KA, Smit A, Riemer C, Bouck J, Gibbs R, Hardison R, Miller W. (2000). PipMaker—a web server for aligning two genomic DNA sequences. Genome Res. 10: 577–586.

38. Tamura K, Stecher G, Peterson D, Filipski A, Kumar S. (2013). MEGA6: Molecular Evolutionary Genetics Analysis version 6.0. Mol Biol Evol. 30: 2725–2729.

39. Kelley LA, Mezulis S, Yates CM, Wass MN, Sternberg MJ. (2015). The Phyre2 web portal for protein modeling, prediction and analysis. Nat Protoc. 10: 845–858.

